# Statistical estimation of sparsity and efficiency for molecular codes

**DOI:** 10.1101/2024.08.13.607773

**Authors:** Jun Hou Fung, Mathieu Carrière, Andrew J. Blumberg

**Affiliations:** Department of Systems Biology, Columbia University, New York, NY, U.S.A.; DataShape, Centre Inria d’Université Côte d’Azur, Sophia Antipolis, France; Irving Institute for Cancer Dynamics and Departments of Mathematics and Computer Science, Columbia University, New York, NY, U.S.A.

## Abstract

A fundamental biological question is to understand how cell types and functions are determined by genomic and proteomic coding. A basic form of this question is to ask if small families of genes or proteins code for cell types. For example, it has been shown that the collection of homeodomain proteins can uniquely delineate all 118 neuron classes in the nematode *C. elegans*. However, unique characterization is neither robust nor rare. Our goal in this paper is to develop a rigorous methodology to characterize molecular codes.

We show that in fact for information-theoretic reasons almost any sufficiently large collection of genes is able to disambiguate cell types, and that this property is not robust to noise. To quantify the discriminative properties of a molecular codebook in a more refined way, we develop new statistics – *partition cardinality* and *partition entropy* – borrowing ideas from coding theory. We prove these are robust to data perturbations, and then apply these in the *C. elegans* example and in cancer. In the worm, we show that the homeodomain transcription factor family is distinguished by coding for cell types sparsely and efficiently compared to a control of randomly selected family of genes. Furthermore, the resolution of cell type identities defined using molecular features increases as the worm embryo develops. In cancer, we perform a pan-cancer study where we use our statistics to quantify interpatient tumor heterogeneity and we identify the chromosome containing the HLA family as sparsely and efficiently coding for melanoma.

## 1. Introduction

A basic question in genomics and proteomics is understanding and describing how cell types and functions are determined by molecular codes. A famous example is the family of homeobox genes, which has long been known to concisely code for complicated phenotypic features in a variety of species.

The nematode *Caenorhabditis elegans* is an ideal organism for studying these questions in systems neuroscience. A mature hermaphrodite worm has a small nervous system comprising of just 302 neurons that fall into 118 neuron classes, which is highly stereotyped across different individuals; to date, it is one of the few animals for which we have essentially a complete atlas of all the different neuron types and the connections between them. Recently, by combining this knowledge with proteomic expression data, it was shown that every neuron class has an unique transcriptomic profile using only genes from the homeodomain transcription factor family [Reilly et al., 2020]. This result was confirmed through further experiments [Leyva-Díaz and Hobert, 2022, Cros and Hobert, 2022, Reilly et al., 2022].

One would hope that this property characterizes this family of genes. However, results of this type are not robust: by removing or corrupting a single gene from the homeobox family, the remaining family may no longer possess the property of being able to uniquely discriminate all neuron classes. This lack of robustness is due to the binary nature of the property: either a neuronal code is able to differentiate all neuronal classes or it is not.

Our goal in this paper is to introduce new statistics that allow us to robustly characterize what is distinctive about molecular codes. Specifically, we propose instead that it is more meaningful to characterize the discriminatory ability of molecular codes on a continuum scale. To this end, we develop novel measures of discriminatory power that are more refined and provably robust to changes in the gene set and bit-flip errors in the codebook. We validate our new statistics by demonstrating that these measures allow us to identify interesting families of genes and proteins that code for cell types.

In this article we make the following contributions:

1. show that unique encoding is a generic property of moderately-sized gene families,
2. introduce two new statistics – *partition cardinality* and *partition entropy* to better quantify the discriminative properties of molecular codebooks,
3. establish mathematically their robustness property and basic behaviour under a null model,
4. use these statistics to analyze homeobox genes as well as other transcription factor families in *C. elegans*,
5. extend the analysis from the L4 larval stage to earlier time points during the worm’s embryonic development, and
6. apply these statistics in a different setting to quantify interpatient tumor heterogeneity in a pan-cancer study.

We regard this article as a first step towards a new kind of coding theory that is relevant to the codes that arise in a biological context.

### Related work

Our work is related to the analysis of combinatorial neural codes in [Curto et al., 2013], which studied co-firing patterns in neural spike train data using a coding theory perspective. We too borrow ideas from coding theory, but here we study molecular codes: our codes consist of genes that are co-expressed in the same cell type or genes that are co-mutated in a tumor, instead of neurons that are functionally active at the same time. In the transcriptomics context, [Casey et al., 2023] have also used information theory to quantify the per-gene expression heterogeneity given a single cell sequencing dataset and devised a clustering algorithm based on entropy. Part of our work explores the converse relationship: we quantify the heterogeneity of a codebook of cell types using genes as features instead. These different examples demonstrate the pervasiveness of biological codes in nature.

## 2. Most families of genes can uniquely delineate all the neuron classes in *C. elegans*

In this section, we show that the homeobox genes are not characterized by their ability to uniquely delineate the neuron classes in the worm. On the contrary, we demonstrate that most random families of sufficient size have this property.

In order to make comparisons beyond homeobox genes, we need to be able to measure the presence or absence of these other genes in each neuron class of *C. elegans*. While [Reilly et al., 2020] produced a dataset based on gold standard protein expression, their experiment extends only to homeodomain proteins, and additional proteomic databases for the worm are mostly incomplete. Therefore, we turn to a reference based on single cell RNA sequencing [Taylor et al., 2021] which provides a more unbiased coverage of other genes in the worm genome.

For 1 ≤ *k* ≤ 100, we repeatedly sampled without replacement the list of genes available in the CeNGEN dataset to obtain 200 collections of size *k*, and then counted the number of collections of genes that were able to uniquely delineate all 128 cell subtypes.

As we can see from Figure 1, most collections with fewer than 25 genes do not possess the unique delineation property. Beyond that, the proportion of collections with the property steadily increases, and on average half the collections of size 45 have the property and more than 90% of the collections of size 65 have the property. Of the 102 genes classified as homeobox in [Reilly et al., 2022], 86 were present in the CeNGEN dataset; a random collection of 86 genes is able to uniquely differentiate all neuron classes 97.5% of the time.

**Figure 1.**
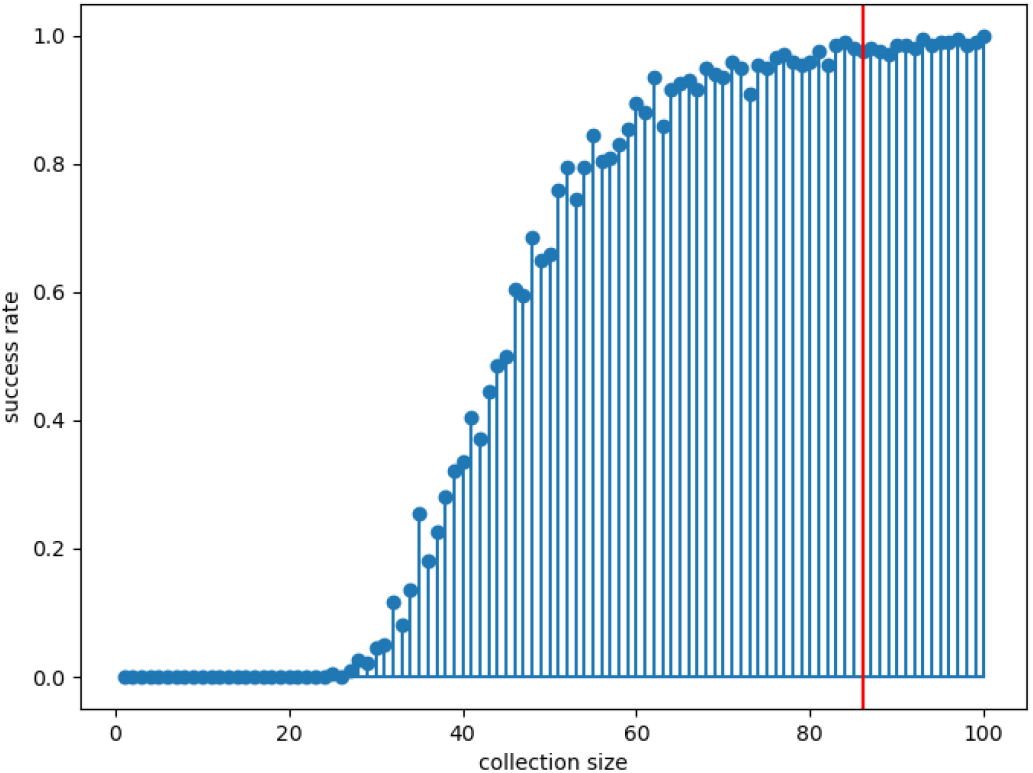
Proportion of random subsets of fixed size with the unique delineation property. Most collections of 50 genes have a probability of *>* 50% of uniquely delineating all the neuron types in *C. elegans*; > 97% of the collections of size 86 (the number of homeobox genes in the reference), shown in red, can uniquely delineate all neuron types.

This analysis leads to two questions we shall address in the remainder of this article. First, can we design a better metric to gauge the discriminatory power of a molecular codebook? Second, if the homeobox genes are not unique in uniquely delineating neuron types, are they still special in some other way, given their biological importance? This latter question reveals our general interest in understanding the distinctive properties of molecular codes.

## 3. Novel metrics robustly quantify the discriminative power of molecular codebooks

We introduce new robust metrics to characterize molecular codes.

### 3.1 Partition cardinality and entropy

We represent the *codebook* of correspondences between neuron class identities and the ensemble of genes or proteins they express as a binary (*N* × *G*)-matrix *E*, where *N* is the set of neurons (or samples more generally), *G* is a set of genes or proteins, and

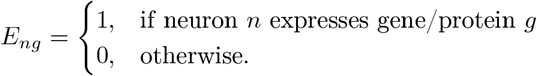

Motivated by our resampling analysis in the previous section, we propose the following statistics to quantify the discriminating power of sets of gene features.

#### Definition 1

Let *E* be a binary (*N* × *G*)-matrix.

i. Given *S* ⊆ *G*, let *n*(*E*_*S*_) be the number of unique rows in the matrix restricted to the columns indicated by *S*.*E*
ii. Let *k* ≥ 1. Define the *k*-th (normalized) *average partition cardinality* of *E* to be

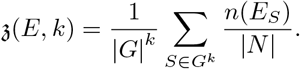

In other words, the submatrix *E*_*S*_ defines a partition of its rows based on grouping together identical rows, *n*(*E*_*S*_) is the cardinality of that partition, and z(*E, k*) is a normalized average of the partition cardinalities over all possible combinations of *k*-elements subsets *S* (drawn with replacement) of columns of *E*.

There is another way to measure the informativeness of a partition, called its *entropy*. Recall, if *P* is a partition of a set of size *N* into parts of size *p*_1_, *p*_2_, …, *p*_*n*_, then the entropy of *P* is given by the formula

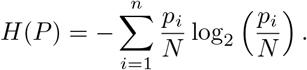

#### Definition 2

Let *E* be a binary (*N* × *G*)-matrix.

i. Given *S* ⊆ *G*, let *H*(*E*_*S*_) denote the entropy of the partition obtained by grouping together the identical rows in the matrix *E* restricted to the columns *S*.
ii. Let *k* ≥ 1. Define the *k*-th (normalized) *average partition entropy* of *E* to be

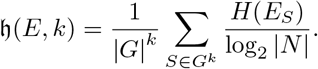

Note that these statistics have been normalized so that 𝔷(*E, k*) ∈ [0, 1] and 𝔥(*E, k*) ∈ [0, 1]. If 𝔷(*E, k*) or 𝔥(*E, k*) is large, meaning that the partitions produced using *k* genes have many blocks or high entropy on average, then the gene family is *efficiently* coding cell identity.

Since both 𝔷(*E, k*) and 𝔥(*E, k*) are obtained by analyzing the partitions of the submatrices *E*_*S*_, it is not surprising that they are related. Specifically, we can show

#### Theorem 3

*Given a binary* (*N* × *G*)*-matrix E, we have for all k* ≥ 1,

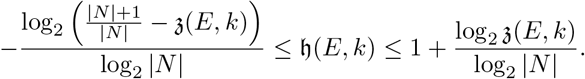

While this suggests that the two statistics should be positively associated, we find they capture different aspects of the codebooks depending on the application. The proof of the theorem, given in the appendix, boils down to the monotonicity of *Rényi entropy* H_*α*_ with respect to the order *α* ∈ [0, ∞]. Rényi entropy is a diversity index that includes as special cases log cardinality H_0_ and Shannon entropy H_1_. Substituting H_*α*_ into our definition above, we can obtain a family of partition statistics that interpolate from (a log variant of) partition cardinality to partition entropy. Intuitively, the order of Rényi entropy is related to the weight placed on larger blocks in the partition, so for example partition entropy is more sensitive to large clusters compared to partition cardinality.

In practice, it is impractical to compute z(*E, k*) and h(*E, k*) exactly: they involve enumerating all size *k* subsets of *G* and constructing partitions in the corresponding restricted matrix. Instead, we can use sample estimators. Given *k*, we randomly generate *b* subsets *S*_1_, *S*_2_, …, *S*_*b*_ ⊆ *G* of size *k*, and define

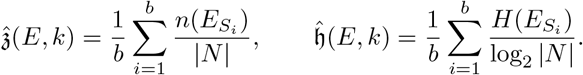

The law of large numbers then implies 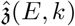 (resp. 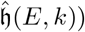) converges to 𝔷(*E, k*) (resp. 𝔥(*E, k*)) almost surely as the number of samples *b* tends to infinity. In the rest of this paper, we use *b* = 300, which is a compromise between computation time and accuracy; by the central limit theorem this corresponds to a theoretical bound on the 95% confidence interval of about *±*5%.

#### Remark 4

If *k* = |*G*|, then these are just the usual bootstrap estimates of the partition cardinality and entropy.

In the applications that follow, the data we have are naturally binarized: a worm neuron is either a particular class or not, so the proteins that comprise its identity program are either translated or not; a patient either has a mutation in a gene or they do not. Even datasets coming from continuous measurements of RNA expression can be analyzed by binarizing into zero or nonzero values, especially as proxies for protein levels: the relationship between RNA measurement and the amount of protein output is complex, but at least we know some nonzero level of the RNA is required for any nonzero protein product.

Nonetheless, there are some cases in which the data we want to analyze are truly continuously distributed. In this case, one simple adaptation is to threshold the data at some level and calculate our statistics on the thresholded values, leading to a family of statistics parametrized by the threshold level. Another option is to fix a clustering method *𝒞*, and define

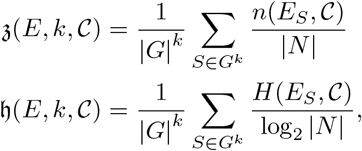

where *n*(*E*_*S*_, *𝒞*) (respectively *H*(*E*_*S*_, *𝒞*)) is the number of clusters (respectively the entropy of the partition) obtained from applying *C* to the rows of *E* restricted to the columns *S*. If *C* is the clustering method that returns clusters consisting of identical observations, then 𝔷(*E, k, 𝔷*) = 𝓏(*E, k*) and 𝔥(*E, k, 𝒞*) = 𝔥(*E, k*), so this is a generalization of partition cardinality and entropy.

### 3.2 Distributions under the null hypothesis

Suppose *E* is a random *N* × *G* Bernoulli matrix, i.e., each entry of the matrix is independently 1 with some probability *p* and 0 with probability 1 − *p*. This serves as a null hypothesis of uninterestingness where there is absolutely no correlation between neurons or genes, so it is useful to study the behaviour of 𝔷(*E, k*) and 𝔥(*E, k*) in this model.

#### Theorem 5

*Let E be a random N* × *G Bernoulli matrix with parameter p. For* 0 *< l* ≤ *k, set π*_*l*_ = *p*^*l*^(1 − *p*)^*k*−*l*^. *Then*,

A. 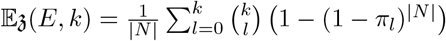
B. 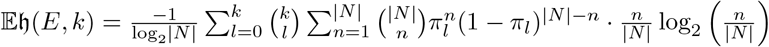

Specializing to the case *p* = 0.5, for large *N* and fixed *k*, we have

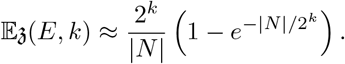

We can use this rough calculation to give a heuristic for selecting the parameter *k*. Specifically, if *k* ≈ log_2_ |*N* |, then 𝔼𝔷(*E, k*) ≈ 1 − *e*^−1^ ≈ 0.63, which is roughly in the middle of the range [0, 1]. Furthermore, if |*N* | = 118, then this heuristic gives *k* ≈ 7. In practice, we find that if *k* is too small then these statistics are not very powerful: if *k* ≪ log_2_ |*N* |, then 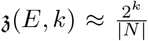 for most *E*. As a compromise, we shall set *k* = 10 for our applications in the worm brain and *k* = 20 in our pan-cancer mutation study.

### 3.3 Wasserstein stability

These statistics also enjoy a important stability property. Regard *E* as a measure on Hamming space *{*0, 1*}*^*N*^ given by

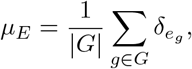

where *e*_*g*_ is the *g*-th column of *E*. If *µ*_*E*_*′* is another measure on *{*0, 1*}*^*N*^ similarly derived from a binary (*N* × *G*^*′*^)-matrix *E*^*′*^, then the *Wasserstein distance* between *E* and *E*^*′*^ is the Wasserstein distance between their corresponding measures, i.e.,

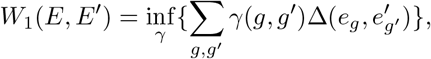

where the infimum is taken over all couplings *γ* : *G×G*^*′*^ → [0, 1], i.e., ∑*g*∈*G γ*(*g, g*,^*′*^) = |*G*^*′*^|^−1^ and 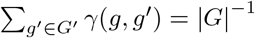 and Δ is the normalized Hamming metric on

Then, we have

#### Theorem 6

*Given E, E*^*′*^ ⊆ *{*0, 1*}*^*N*^, *for k* ≥ 1, *we have*

a. |𝔷(*E, k*) − 𝔷(*E*^*′*^, *k*)| ≤ *k · W*_1_(*E, E*^*′*^)
b. |𝔥(*E, k*) − 𝔥(*E*^*′*^, *k*)| ≤ *k*(1 + (log |*N* |)^−1^) *· W*_1_(*E, E*^*′*^).

This stability result guarantees that our statistics are robust to minor changes in molecular features and to errors in the codebook. Hence, unlike simply asking whether a given molecular code can uniquely delineate all cell types, partition cardinality and entropy metrics are suitable for the analysis of noisy or corrupted data.

## 4. A tradeoff between coding efficiency and sparsity

Transcriptomic codes can have quite variable sparsity: for instance, in the CeNGEN database, an average homeobox gene is only expressed in 22% of neurons compared to a global gene average of 41%. From theorem 5, we can deduce that in the Bernoulli model a smaller *p* leads to a decrease in 𝔼𝔷(*E, k*) and 𝔼𝔥(*E, k*). This can be explained by the fact that the sparsity constraint yields more correlated codes, and hence the expected coding efficiency to differentiate between different neurons is reduced.

However, in a real dataset, the level of sparsity is not given and has to be inferred. Given a binary matrix *E*, we define its *effective sparsity* to be the proportion of its entries that are zeroes or the proportion of its entries that are ones, whichever is the larger. By definition, the effective sparsity takes values between 0.5 and 1: an effective sparsity of 0.5 means that the two binary classes are perfectly balanced, while an effective sparsity of 1 means that the matrix is either all zeros or all ones. Next, we conducted a simulation study using the Bernoulli null model introduced in the previous section. First we randomly and uniformly selected *p* ∈ [0, 1] and generated a random Bernoulli (100 × *k*)-matrix *E* with parameter *p*. Then, we computed the partition cardinality 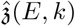 and entropy 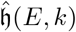 for *k* = 10. We repeated this process 1000 times, and plotted the results against the effective sparsity in Figure 2. As expected, as sparsity increases, partition cardinality and partition entropy decreases reflecting a lower diversity of codes. This result implies that the values of partition cardinality and entropy should be interpreted in the context of the dataset sparsity.

**Figure 2.**
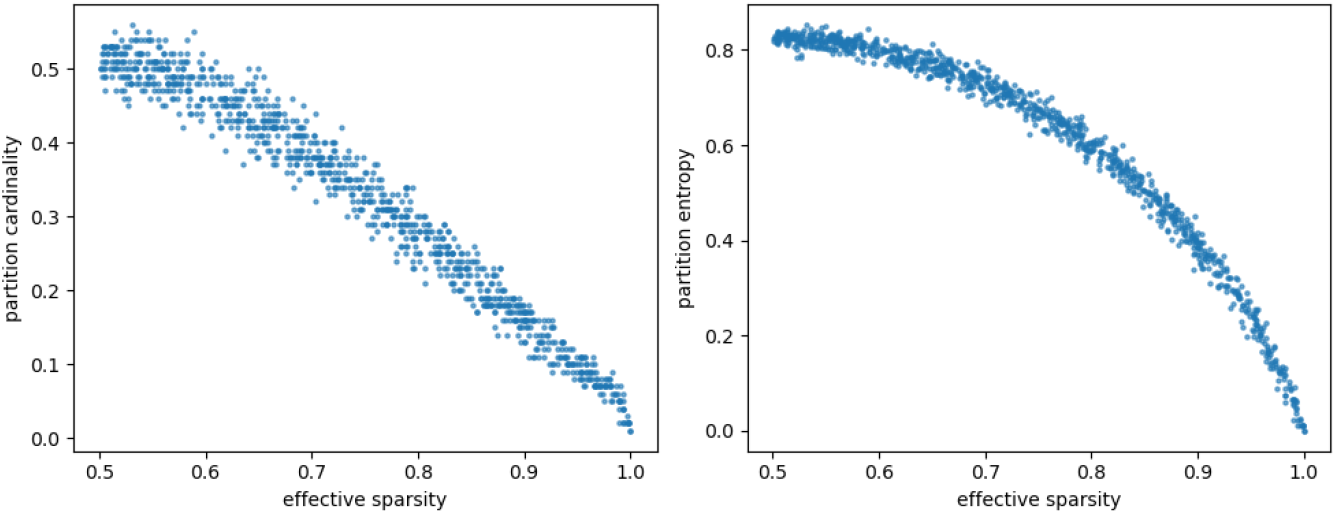
Partition cardinalities and partition entropies decrease with increasing effective sparsity (the size of the majority binary class) in a random Bernoulli model, reflecting a tradeoff between the discriminative ability and the code sparsity.

### Remark 7

Our notion of effective sparsity does not capture correlations between cells. However, biologically speaking our simple measurement of sparsity may be sufficient and more relevant: to produce a gene or protein a cell has to recruit the appropriate resources and transcriptional and translational machinery. In this simplistic model, a cell incurs an energetic cost in order to express a wide variety of diverse genes. Hence, there is a tradeoff between expressing many different genes (low effective sparsity) and generating an efficient code that distinguishes cell types.

## 5. Applications to *C. elegans* neuronal codes

### 5.1 Partition cardinality and entropy are stable between codebook versions

[Reilly et al., 2020, Supplementary Table 2] provides a codebook *E* consisting of |*N* | = 118 neuron classes and |*G*| = 101 homeodomain proteins. We calculated the values of 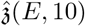 and 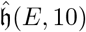, and visualized the results in Figure 3.

**Figure 3.**
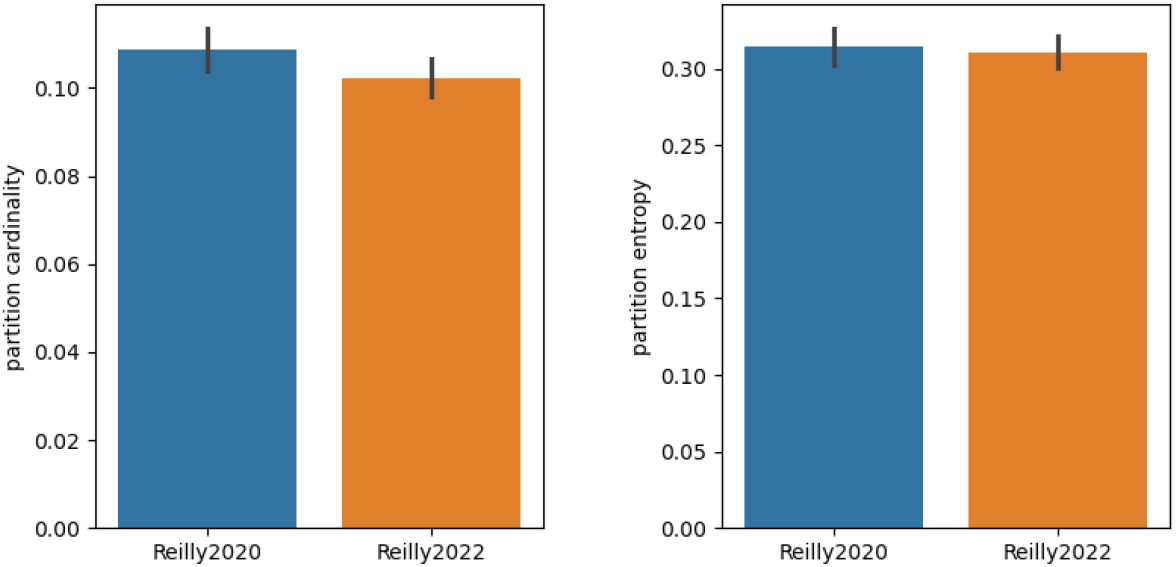
Values 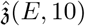 and 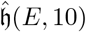 for the Reilly et al. datasets [Reilly et al., 2020, 2022]. There is no statistically significant difference between the values of the either the partition cardinality (two-sample Welch *t*-test: *t* = 1.933, *p* = 0.054) or partition entropy (two-sample Welch *t*-test: *t* = 0.473, *p* = 0.636) obtained from the two versions of the *C. elegans* homeodomain protein codebook.

We note that an updated expression matrix is available from [Reilly et al., 2022, S1 Table]. On the one hand, it includes the remaining homeobox gene *ceh-84* [Reilly and Hobert, 2020] (which is not detected anywhere in the mature worm nervous system). On the other hand, it divides some but not all of the 118 neuron classes into further subtypes for a new total of 128 classes. Furthermore, according to the updated table homeobox genes do *not* uniquely delineate neuronal identities: the neurons URA and URB now have the same code. Despite these differences, our partition cardinality and entropy statistics do not change significantly, in line with expectations from our stability theorem. This kind of technical variability in data precisely demonstrates the need for robust statistics for analysis.

### 5.2 The homeodomain and nuclear receptor transcriptional codes sparsely delineate neuron classes

We now apply our framework to study the transcriptional codes provided in the CeNGEN dataset [Taylor et al., 2021]. In addition to homeobox genes, we analyze more broadly genes from other transcription factor families. In *C. elegans*, almost 1,000 transcription factors (TFs) have been identified [Fuxman Bass et al., 2016]. In order to perform their function, TFs need to be able to bind to DNA, and consequently they possess a DNA binding domain, and we can classify TFs by their DNA binding domains. One example of a TF family is the homeodomain (HD) family we examined above. Other families include the zinc finger (ZF) family, the nuclear hormone receptor (NHR) family, the basic helix-loop-helix (bHLH) family, and the basic leucine zipper (bZIP) family.

We stratified the available TFs in the CeNGEN dataset by TF family by crossreferencing with the table provided in Fuxman Bass et al. [Fuxman Bass et al., 2016, Dataset EV3]. We recovered 86/101 HD genes, 415/565 ZF genes, 189/268 NHR genes, 24/41 bHLH genes, and 25/32 bZIP genes. Then, we restricted the CeNGEN expression matrix to these gene subsets and inferred the partition cardinality and entropy of these families. We also computed the effective sparsity of each of these TF families. For control, we subsampled the entire CeNGEN dataset to obtain 5000 collections of 86 genes each, and also computed the sparsity and partition statistics of these random collections. The results are shown in Figure 4.

**Figure 4.**
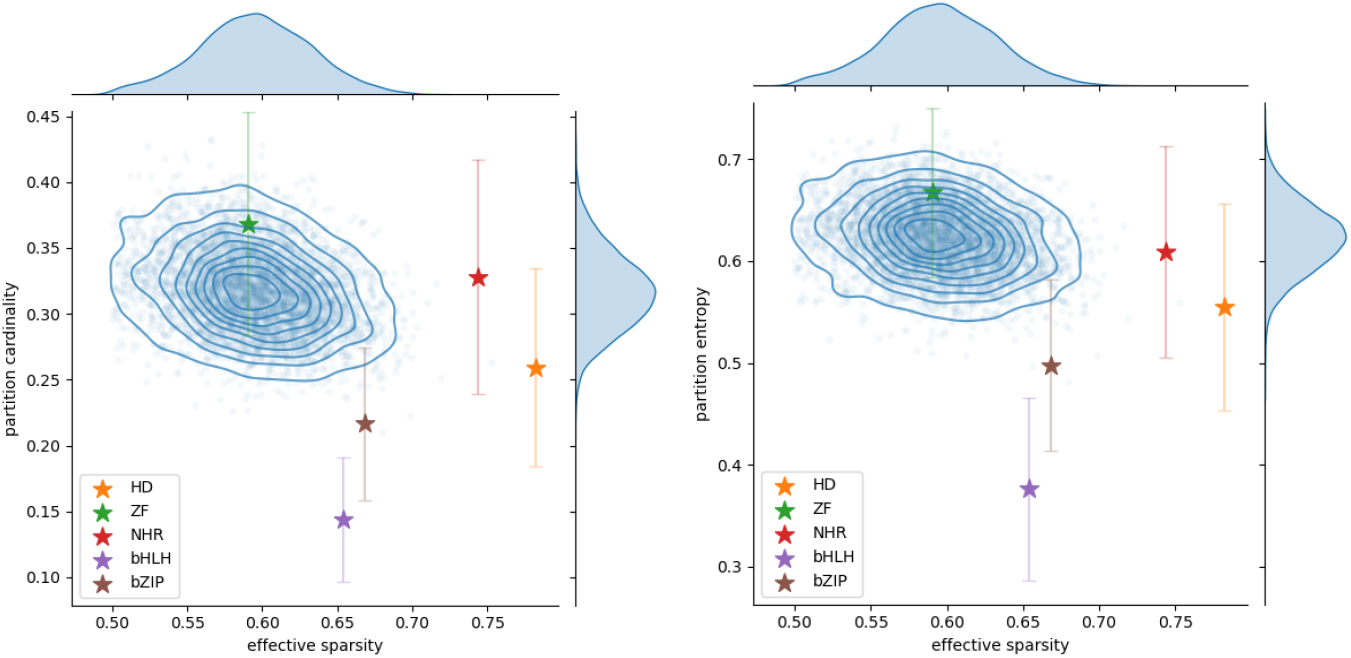
Sparsity, partition cardinality and partition entropy of the *C. elegans* transcriptomic code, specialized to various transcription factor families. All of the transcription factor families, except the zinc-finger family, have distinct sparsity levels and statistics compared to a random gene family (in blue). More specifically, the homeodomain and nuclear hormone receptor families have much higher effective sparsity, but still maintain a relatively high partition cardinality and entropy.

We see firstly that out of the five TF families considered, four of them are sparser than average, the exception being the largest zinc finger family. In particular, the homeodomain and nuclear hormone receptor families are significantly sparser than a random collection of genes. Furthermore, we see the bHLH and bZIP families have lower partition cardinality and entropy compared with the random families. This is consistent with the observation that in fact these families do not uniquely delineate all the neuron classes: the entire bHLH and bZIP families could only differentiate 74 and 104 classes, respectively, out of 128. Finally, we see that despite being much sparser, the homeodomain and nuclear receptor families still have comparatively high partition cardinality and entropy.

To further investigate this last point, we repeated our subsampling experiment but only with the genes with sparsity no greater than 0.3. Out of 1000 such collections generated, only 21 had greater partition cardinality than the homeodomain family (*p* = 0.021) and only 1 had greater partition entropy (*p* = 0.001). As for the NHR family, none had greater partition cardinality or entropy (*p <* 0.001). Altogether, this shows that not only do the homeodomain and nuclear hormone receptor transcriptional codes uniquely delineate neuron classes, but they do so economically.

### 5.3 Molecular disambiguation of cell type identities increases with embryo time

Neuronal identities are gradually acquired during embryonic development. Therefore, it is interesting to analyze how the ability of transcriptomic codebooks to differentiate neuron types changes over the course of development. [Packer et al., 2019] performed single-cell sequencing of over 80,000 cells from *C. elegans* embryos from gastrulation to the cuticle synthesis (pre-larval stages). The data is binned into nine time intervals by inferred embryo times, and then aggregated by approximate cell types. For each time bin, we get a cell type-gene expression matrix. Note both the number of distinct cell types and the cell type annotations change over time; the number of cell types in each period is summarized in table 1.

We computed the partition cardinality and entropy for each time bin separately. As before, we also broke down the gene features by TF family. The results are shown in Figure 5. Excepting the first time bin (embryo time 170–210 minutes) which is anomalous due to extremely low number of distinct cell types, we observe that there is a tendency for the partition cardinality and entropy to increase during the later stages of larval development. This is consistent with the notion that many of these TFs, especially those in the homeodomain family, exert their greatest effects in neuronal identity specification as the worm finishes maturing due to their ongoing roles in the adult worm. We also see in this dataset that the homeobox genes have significantly higher partition cardinality and entropy compared to a random gene ensemble as well as to other TF families, confirming their specific role in efficiently coding neuronal identities.

**Table 1.** Number of distinct cell types in each time interval from the [Packer et al., 2019] dataset.

| Time bin (mins.) | Number of cell types |
| --- | --- |
| 170–210 | 11 |
| 210–270 | 54 |
| 270–330 | 101 |
| 330–390 | 126 |
| 390–450 | 113 |
| 450–510 | 106 |
| 510–580 | 99 |
| 580–650 | 87 |
| > 650 | 62 |

**Figure 5.**
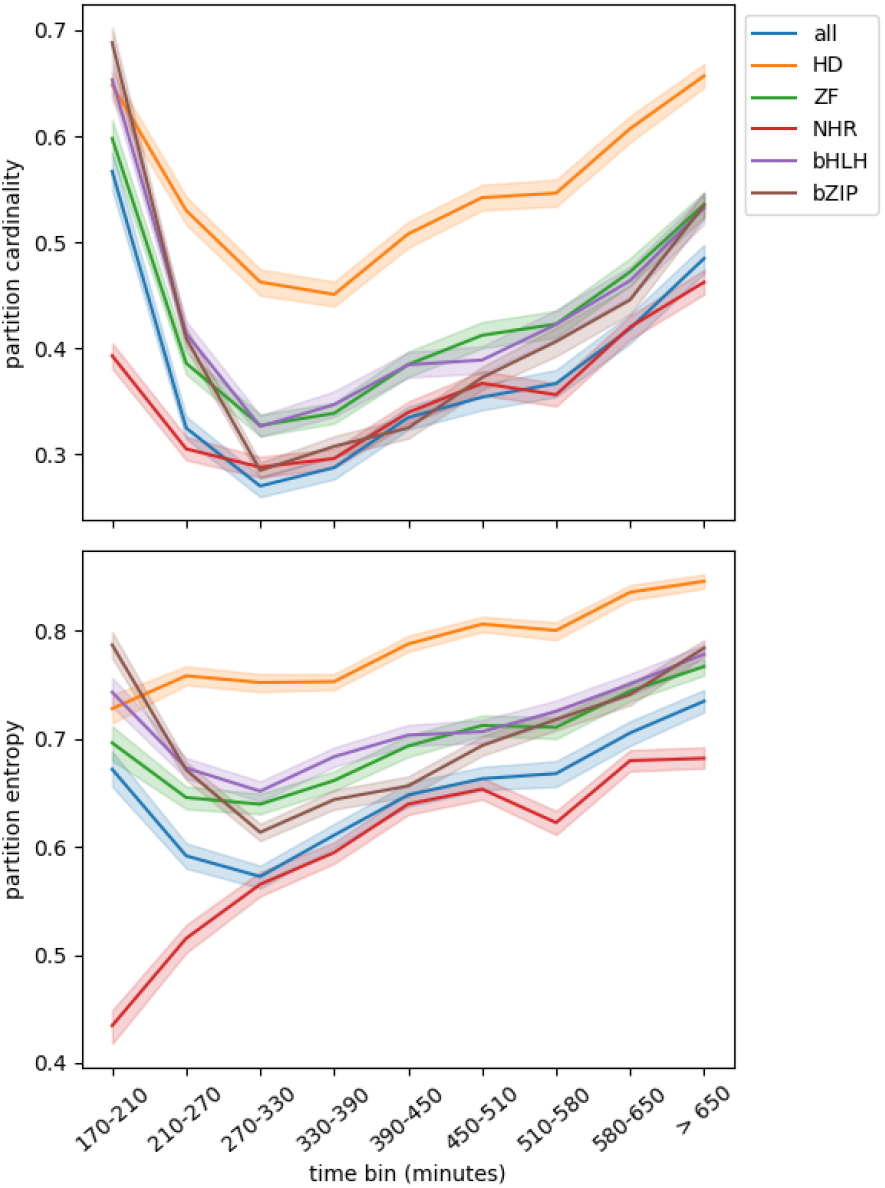
Mean and standard error (shaded) of the partition cardinality and partition entropy computed on a lineage-resolved molecular atlas [Packer et al., 2019], grouped by transcription factor family. There is a general trend of increasing partition cardinality and entropy with maturation time across all transcription factor families. The homeodomain family have the highest partition cardinalities and entropies compared to the other transcription factor families and to all genes.

## 6. Somatic mutation profile diversity in cancers

We further demonstrate the versatility of our method by studying somatic mutational codes in cancers. Tumor heterogeneity is an important consideration in cancer biology; in particular, *interpatient* heterogeneity presents profound challenges for therapy: the goal of personalized and precision medicine is to overcome this heterogeneity by tailoring treatments to the specific molecular and phenotypic features of each cancer. It is therefore crucial to quantify interpatient tumor heterogeneity when devising new treatments [Kashyap et al., 2022].

One way in which tumors vary between patients is through the genetic mutations they harbor, which gives insights into their etiology and pathways of disease progression. We conducted a pan-cancer study using targeted screen mutational data from the COSMIC database [Tate et al., 2018]. We grouped together samples into cancer types by combining primary site and primary histology information and selected 42 cancer types each with at least 1000 tumor samples present. By subsetting the list of 743 cancer census genes to those that are mutated in at least 0.1% of the samples for each cancer type separately, we can encode each sample by its mutational profile. Basic frequencies and co-occurrences of various alterations are commonly studied, but using partition statistics we can also analyze the higher-order diversity of these mutational profiles. For example, a low partition cardinality or entropy means that the mutational codes are not able to separate different samples: in other words, the samples are genomically homogeneous. On the other hand, a high partition cardinality or entropy means that there is high tumor heterogeneity across samples. To speed up calculations we repeatedly generated subsamples of 1000 tumors 100 times prior to computing these statistics for each cancer type. Figure 6 shows the distribution of these statistics for each cancer type. We see interpatient heterogeneity varies widely across cancer types. Hematopoietic and lymphoid neoplasms (e.g., acute myeloid leukemias) and breast carcinomas have the highest partition cardinality while salivary gland carcinomas and biliary tract carcinomas have the highest partition entropy.

**Figure 6.**
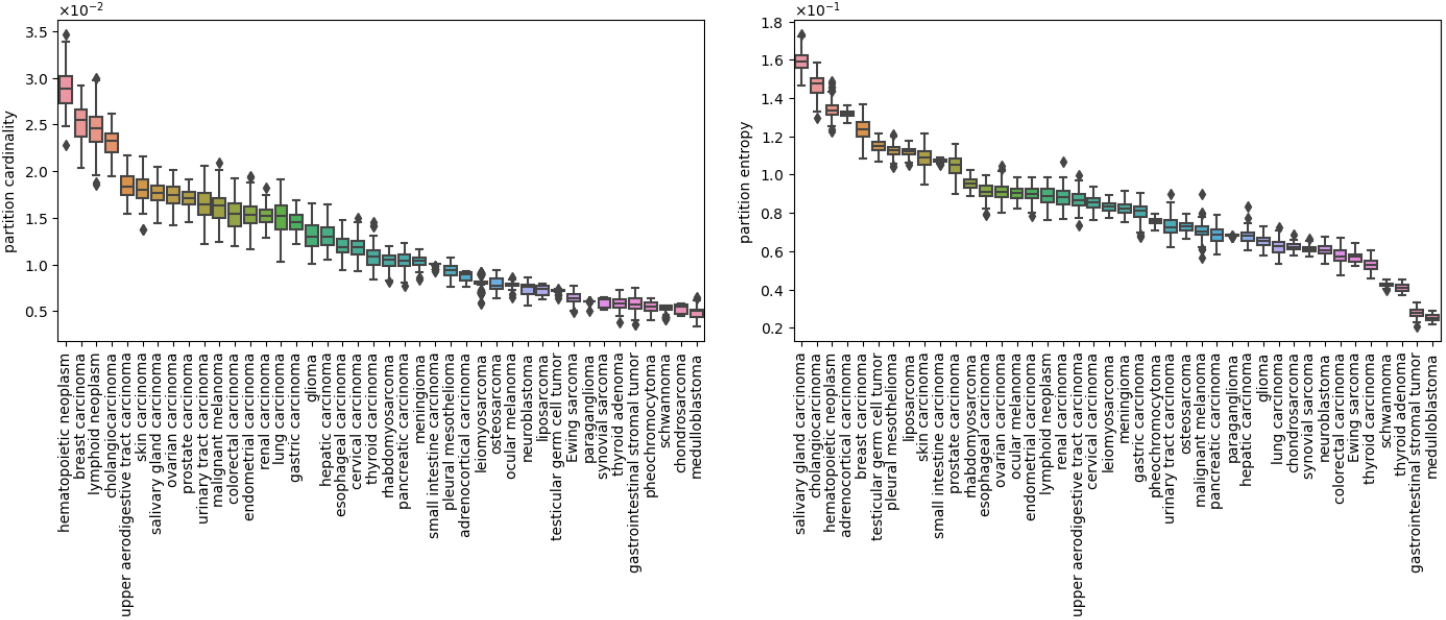
Pan-cancer interpatient heterogeneity as measured by partition cardinality and entropy. Both statistics identify hematopoietic neoplasms, breast carcinomas, cholangiocarcinomas as having the highest diversity, while schwannomas, thyroid adenomas, gastrointestinal stromal tumors and medulloblastomas have the lowest heterogeneity.

In addition to studying interpatient heterogeneity, we can also zoom in on specific cancer types and examine the per-chromosome heterogeneity to identify the genomic regions that are most diverse. We decided to focus on malignant melanomas and gliomas. We restricted the same COSMIC dataset to 1898 gliomas and 3597 melanomas each with at least 10 mutated cancer census genes. For each cohort we separately filtered out uncommonly mutated genes that are not mutated in at least 5 samples. Then, we computed the sparsities and the partition cardinalities and entropies of the melanoma and glioma mutational codes, stratified by chromosome. As controls, we also computed these statistics for 100 random gene collections of size 20. The results are shown in Figure 7. In some cases, the variations in partition cardinality and entropy are likely due to sparsity differences. However, there are also exceptions to this rule. For instance, the chromosome 6 code is the sparsest in the melanoma cohort but surprisingly has one of the highest partition cardinality. Chromosome 6 contains the HLA locus which is one of the most polymorphic regions in the human genome, so it is interesting to find that cancer mutations in this chromosome are likewise especially diverse. In the glioma cohort, we see that the chromosome 4 code is one of the least sparse, but it has a high partition entropy relative even to other codes of similar sparsity. Other differences between chromosomes have more mundane explanations: for instance, as the smallest human autosome, chromosome 21 has a code that often appears as an outlier due to its size.

**Figure 7.**
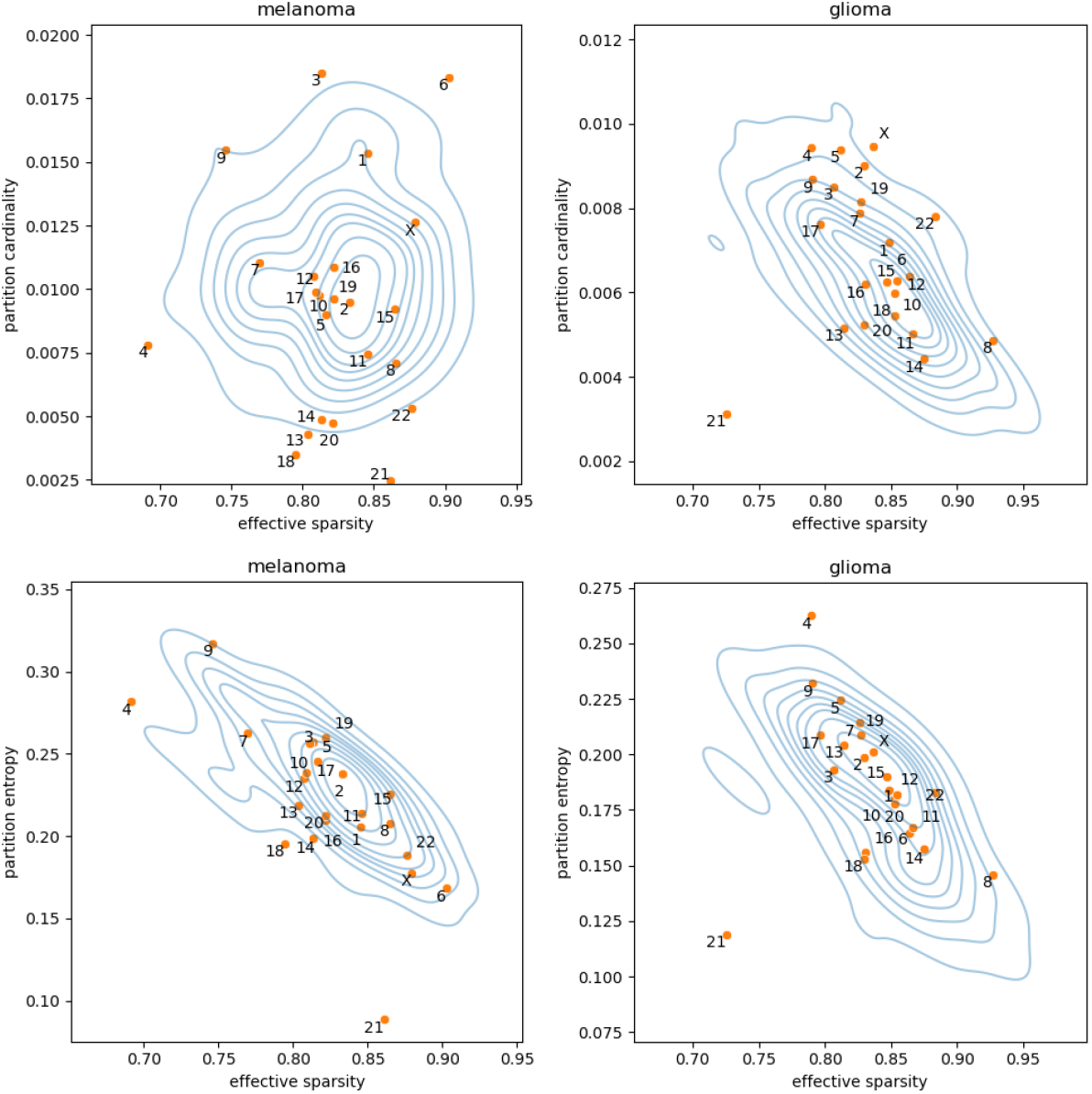
The partition cardinalities and entropies of mutational codes of gliomas and melanomas stratified by chromosome. Partition cardinality analysis identifies chromosome 6 as simultaneously having high diversity and high sparsity in melanomas.

## 7. Discussion

In this article, we sought to unravel the combinatorial logic behind molecular codes (e.g., genomics and proteomics) and cell identity specification by robustly quantifying the joint discriminative power and redundancy of various families of genes.

We were originally motivated by gene expression and neuronal identity specification in *C. elegans*. In recent decades, a key organizing principle for nervous system development has emerged based around the concept of “terminal selectors” [Hobert, 2008]. A terminal selector is a gene or protein that initiates and then continuously maintains the terminally differentiated cell states, thereby defining the identity of mature cells. Studies in *C. elegans* have shown that these terminal selectors are used – and reused – in complex combinations to ultimately determine neuron-type identity alongside parallel regulatory routines.

Many known terminal selectors in *C. elegans* are homeodomain transcription factors; furthermore, [Reilly et al., 2020] had shown the expression of these homeobox genes allow for a unique decoding of neuron type. However, this may simply be a combinatorial, and not biological, consequence: we showed that this property is shared by most families of genes by chance, terminal selectors or not. Furthermore, we expect that uniqueness alone is typically insufficient for meaningful biological interpretation, since it is a fragile property that may be impacted by a small number of mutations. In order to disambiguate homeodomain genes from other collections of genes and to test the fragility of the transcriptomic code, we introduced robust statistics and found that the codebooks provided by the homeobox genes are in fact no more efficient than a random collection of genes at encoding neuronal identity, suggesting some other objective may be in play. We hypothesized that this phenomenon is the result of a tradeoff between efficiency and sparsity. There are over 20,000 genes in the *C. elegans* genome, and expressing each gene adds to the burden both in terms of the transcriptional and translational machinery needed to maintain and regulate it, so there is a biological rationale to express only a smaller number of highly specific genes within each cell type. After adjusting for sparsity, we find that homeobox genes are indeed better at encoding the diversity of neuron types in the worm. Furthermore, we also found evidence that this discriminative power increases with embryonic development. Misexpression studies have revealed that the activity of terminal selectors can be modulated based on specific developmental timings and it would be interesting to explore whether this can be reproduced using our metrics of code diversity.

While our most striking applications were to molecular codebooks for cell type identity, our methods are also relevant for studying other kinds of codes. For example, we used our statistics to study the mutational codes that characterize tumor samples and to devise measures of interpatient heterogeneity that are global and potentially capture different information from simply looking at frequencies of known alterations implicated in cancer pathways. Here, measures indicating high levels of sparsity and expressiveness imply heterogeneous cancers and therefore suggest treatment protocols must be adapted to the specific patient. This work is in comparatively preliminary stages, and we regard these results as a suggestive proof of concept. Work in progress compares our novel and robust measures of heterogeneity with other known measures of genomic variability in cancers.

We believe that this paper is a first step towards a theory of codes that is adapted to the properties of biological codes. Notably, one lesson that we can draw from our study is while mathematical coding theory seems to be an obvious tool to study biological codes, the usual concepts from that field, such as error control and efficiency, are insufficient to characterize the properties of these codes. For instance, [Curto et al., 2013] suggested that receptive field codes struck a compromise between error correction and encoding stimuli relationships, in parallel with the compromise we discovered in molecular codes between coding efficiency and sparsity. Another lesson towards applying coding-theoretic analyses is that there is a need to develop summaries of these various sorts of codes that are robust to small perturbations, perhaps by using an ensemble average approach as we have done here. Finally, many biological systems are often thought of as layers of interlocking networks; for example, nervous systems have vertices given by neurons (or neural regions) and edges given by synaptic, gap junctional, and neuropeptidergic signalling connectomes and gene variants exert tumorigenic effects not in isolation but through cascades in gene regulatory networks. Here, we studied how the *vertices* of such graphs are encoded molecularly; a natural follow-up would be to understand how and to what extent the *edges* and other higher-order interactions are also encoded in gene expression profiles.

## Appendix A. The unique coding of homeobox genes is not robust

Analyzing the data from [Reilly et al., 2020, Supplementary Table 2], we observe

- If the gene *lin-39* is omitted, then the neurons FLP and PVD have the same code.
- If the gene *zag-1* is omitted, then the neurons RMH and SMD have the same code.
- If the gene *ceh-43* is omitted, then the neurons URA and URB have the same code.

Note that however the neurons FLP and PVD, despite being physically distant in the adult worm (FLP in the head and PVD in the body), have similar anatomical structures and play similar roles in nociception and thermosensation.

## Appendix

### B. Proofs

#### B.1 Proof of theorem 3

The relationship between entropy and cardinality is well-exposed Radhakrishnan [2003]. Our bounds between the partition cardinality and partition entropy of a codebook are straightforwardly derived from the usual bounds between regular cardinality and entropy. Let *P* = (*p*_1_, …, *p*_*n*_) be a partition of |*N* | elements into *n*(*P*) = *n* inhabited parts, and write *H*(*P*) for the entropy of the partition *P*.

##### Lemma 8

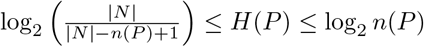

*Proof*. The right inequality is well-known: the maximum entropy of a random distribution is achieved by a uniform distribution, and with *n* outcomes this maximum entropy is equal to log_2_ *n*.

The left inequality follows from the fact that (Shannon) entropy is bounded below by the *min-entropy*, which is defined as 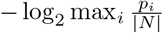 With no constraints, the maximum *p*_*i*_ could be |*N* |, in which case the min-entropy is zero. However, we require that each part has positive mass, i.e, *p*_*i*_ ≥ 1 for each *i* = 1, …, *n*. This means that max_*i*_ 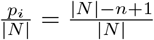, which yields the desired inequality.

##### Remark 9

The left inequality in the lemma above can be made tighter: we have

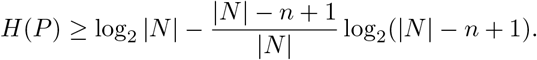

We can now prove the theorem:

**Theorem**. *Given a binary* (*N* × *G*)*-matrix E, we have for all k* ≥ 1,

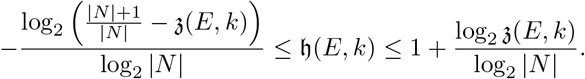

*Proof*. We prove the right inequality: plugging the bound from lemma 8 into the definition of 𝔥(*E, k*), we have

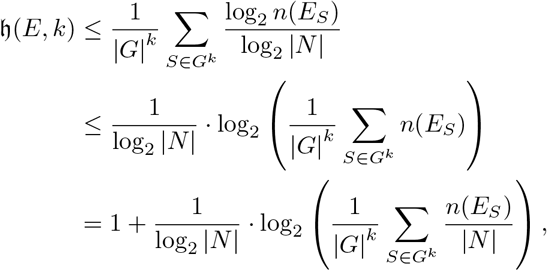

where we have used Jensen’s inequality in the second line.

The proof of the left inequality follows from similar manipulations, using the other bound from lemma 8.

#### B.2 Proof of theorem 5

We restate the theorem.

**Theorem**. *Let E be a random N* × *G Bernoulli matrix with parameter p. For* 0 *< l* ≤ *k, set π*_*l*_ = *p*^*l*^(1 − *p*)^*k*−*l*^. *Then*,

a. 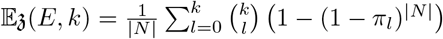
b. 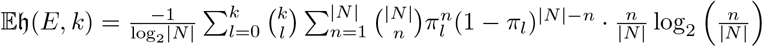

*Proof of (a)*. Since *E* is a random Bernoulli matrix, 𝔼*n*(*E*_*S*_) =: 𝔼*n*(*E*_*k*_) depends only on *k* = |*S*|. So, by linearity of expectation,

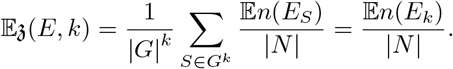

For a string *v* ∈ {0, 1}^*k*^, let 1_*v*_ be the indicator random variable that takes value 1 if *v* appears at least once as one of the |*N* | rows in *E*_*k*_, and is zero otherwise. Then, 1_*v*_ is a Bernoulli random variable with success probability 1 − (1 − *p*^*l*^(1 − *p*)^*k*−*l*^)^|*N*|^, where *l* = *l*(*v*) is the number of 1’s in the string *v*.

Note that

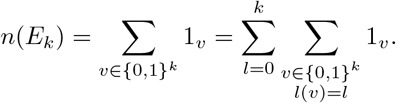

Hence,

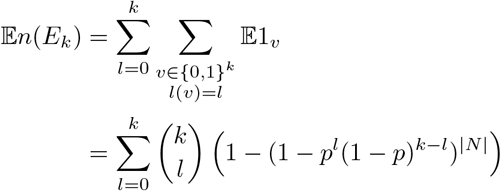

as desired.

*Proof of (b)*. Analogously to part (a), we have

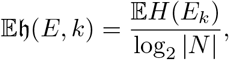

Where

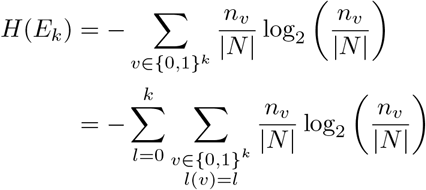

and *n*_*v*_ is the number of rows of a |*N* | × *k* random Bernoulli matrix labelled *v* ∈ {0, 1}^*k*^. Note that *n*_*v*_ ∼ Bin(|*N* |, *π*_*l*(*v*)_) is binomially distributed. Hence,

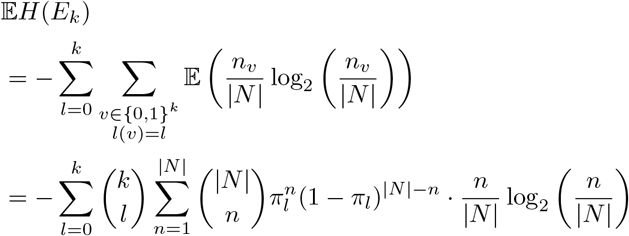

#### B.3 Proof of theorem 6

We prove a generalized version of theorem 6, which asserts that the partition cardinality and entropy we defined in the main text are robust with respect to minor changes measured using the Wasserstein distance. First, we establish a Lipschitz bound on the partition cardinality function.

##### Lemma 10

*Let f* : ({0, 1}^*N*^)^*k*^ → [0, 1] *be the function* 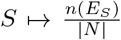. *Then f is* 1*-Lipschitz with respect to the sum metric on its domain*.

*Proof*. Since ({0, 1}^*N*^)^*k*^ has the sum metric, we can work coordinate-by-coordinate, i.e., it suffices to show that

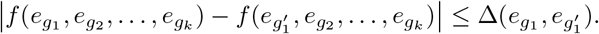

Since *E*_*{g*1_, *g*_2_, …,*g*_*k*_ } and 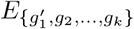 differ in o nly 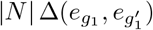 rows, we conclude that *n*(*E*_*{g*1_, *g*_2_, …,*g*_*k*_ }) and 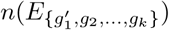 can differ b y a t m ost 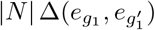. Therefore,

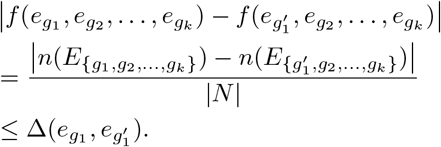

Now, given any measure *µ* on {0, 1}^*N*^, let us define

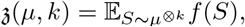

where *µ*^⊗*k*^ is the product measure on ({0, 1}^*N*^)^*k*^. Note that this definition generalizes the partition cardinality function from above: if *E* = *{e*_*g*_}_*g*∈*G*_ ⊆ {0, 1}^*N*^, then we have 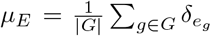, and 𝔷(*µ*_*E*_, *k*) = 𝔷(*E, k*). Furthermore, by definition *W*_1_(*E, E*^*′*^) = *W*_1_(*µ*_*E*_, *µ*_*E*_*′*).

We are now ready to prove the first part of theorem 6.

##### Theorem 11

*Let µ, ν be two measures on {*0, 1}^*N*^. *Then*,

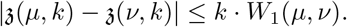

*Proof*. Given measures *µ, ν* on {0, 1}^*N*^, we have

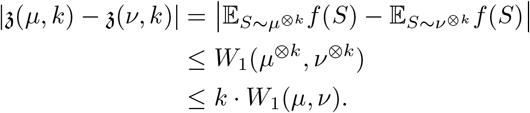

The second line follows from lemma 10 and Kantorovich-Rubinstein duality [Villani, 2008].

The proof of the second part of (6) is similar, once we establish a corresponding Lipschitz bound.

##### Lemma 12

*Let h* : ({0, 1}^*N*^)^*k*^ → [0, 1] *be the function* 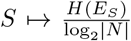. *Then h is* (1 + (log2 |*N* |)^−1^)*-Lipschitz*.

*Proof*. As before, we just need to show

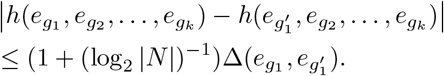

The maximum entropy difference between *E*_*{g*1_, *g*_2_, …,*g*_*k*_ } and 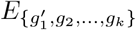 is achieved when one of the partitions is obtained from the other by splitting 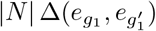 singletons into their own individual clusters. Extracting a singleton from a partition of size *p* increases the entropy by

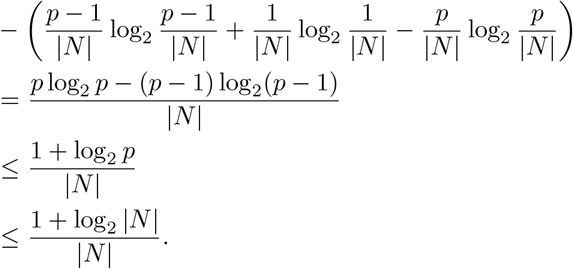

Consequently,

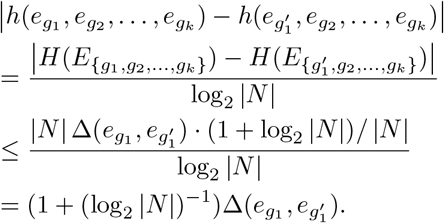

Now, define

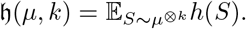

This generalizes the partition entropy function: we have h(*µ*_*E*_, *k*) = h(*E, k*). We can prove the second part of theorem 6.

##### Theorem 13

*Let µ, ν be two measures on {*0, 1}^*N*^. *Then*,

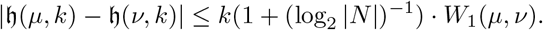

*Proof*. Same proof as theorem 11, except using lemma 12 instead.

## Availability of data and materials

No new data were generated or analyzed in support of this research. The preprocessed binary *C. elegans* homeodomain protein expression tables were obtained from [Reilly et al., 2020, Supplementary Table 2] and [Reilly et al., 2022, Supplementary Table 1]. The binary gene expression table for L4 stage *C. elegans* is available from [Taylor et al., 2021], at https://www.cengen.org; we used the “2 (medium)”-thresholded table. The single cell RNA-sequencing data for *C. elegans* during embryogenesis was retrieved from the Gene Expression Omnibus under accession code GSE126954. The human somatic mutation dataset was retrieved from COSMIC [Tate et al., 2018], at https://cancer.sanger.ac.uk/cosmic; we used the “Complete Targeted Screens Mutant v99 GRCh38” table.

All code and scripts used for the analyses are stored on GitHub, at https://github.com/jhfung/RobustCodeStats.

## Author contributions statement

J.H.F. conceived and conducted the analysis and analyzed the results. J.H.F. wrote and reviewed the manuscript. A.J.B. and M.C. helped conceptualize the work and reviewed and edited the manuscript.

## Acknowledgments

We thank Raul Rabadan and Eviatar Yemini for helpful conversations about this project.

This work is supported in part by funds from the National Science Foundation (NSF: #1912194). Blumberg was supported in part by NSF grant DMS-2311338 and ONR grant N00014-22-1-2679.

